# Nanoscale profiling of evolving intermolecular interactions in ageing FUS condensates

**DOI:** 10.1101/2023.12.21.572955

**Authors:** Alyssa Miller, Zenon Toprakcioglu, Seema Qamar, Peter St. George-Hyslop, F. Simone Ruggeri, Tuomas P. J. Knowles, Michele Vendruscolo

## Abstract

In addition to the native state, proteins can form liquid-like condensates, viscoelastic condensates, such as gels, as well as solid-like condensates, such as amyloid fibrils, crystals and amorphous materials. The material properties of these condensates play important roles in their cellular functions, with aberrant liquid-to-solid phase transitions having been implicated in neurodegenerative diseases. However, the molecular changes and resultant material properties across the whole phase space of condensates are complex and yet to be fully understood. The extreme sensitivity to their environment, which enables their biological function, is also what makes protein condensates particularly challenging experimental targets. Here, we provide a characterisation of the ageing behaviour of the full-length fused in sarcoma (FUS) protein. We achieve this goal by using a microfluidic sample deposition technology to enable the application of surface-based techniques to the study of biological condensates. We first demonstrate that we maintain relevant structural features of condensates in physiologically-relevant conditions on surfaces. Then, using a combination of atomic force microscopy and vibrational spectroscopy, we characterise the spatio-temporal changes in the structure and mechanical properties of the condensates to reveal local phase transitions in individual condensates. We observe that initially dynamic, fluid-like condensates undergo a global increase in elastic response conferred by an increase in the density of cation-π intermolecular interactions. Solid-like structures form first at condensate-solvent interfaces, before heterogeneously propagating throughout the aged fluid core. These solid structures are composed of heterogenous, non-amyloid β-sheets, which are stabilised by hydrogen-bonding interactions not observed in the fluid state. Overall, this study identifies the molecular conformations associated with different physical states of FUS condensates, establishing a technology platform to understand the role of phase behaviour in condensate function and dysfunction.

## Introduction

Phase transitions of proteins have long been the subject of interest in molecular biology, from liquid to solid transitions associated with amyloid formation (*1*), to liquid crystal formation in crystallographic studies (*2*). Recently, protein phase transitions have been reported in cell models via the formation of biomolecular condensates (*3*, *4*).

A protein whose function has been linked to phase separation is the RNA-binding protein fused in sarcoma (FUS) (*5–7*). The ability of FUS to form dynamic, fluid-like assemblies plays an important role in RNA metabolism and DNA damage repair processes (*5–7*). This type of behaviour is imparted by certain sequence features, including a long, intrinsically disordered region, which confers high levels of conformational freedom (*8*), and enrichment in particularly amino acids, especially charged amino acids (*9*, *10*). However, FUS exhibits further phase transitions characterised by the loss of fluid-like behaviour, which have been linked with amyotrophic lateral sclerosis (ALS) (*11*). Evidence is emerging that these phase transitions occur heterogeneously within condesates, with solid formation occurring first at condesate-solvent interfaces (*12*, *13*). ALS-associated mutations have been shown to dysregulate these phase changes, resulting in the formation of large, insoluble assemblies, suggesting a possible pathogenic role of the liquid-to-solid transition (*11*).

Given these observations, it is desirable to understand how the different material states of FUS condensates relate to their physiological and pathological behaviours. More generally, as diverse material properties have been described for biomolecular condensates, it is becoming increasingly clear that material properties play an important role in their functions in health and disease (*14–16*). However, the molecular changes and resultant material properties across the whole phase space of protein condensates, including those formed by FUS, are complex and yet to be fully elucidated.

Efforts to understand the complex changes associated with protein condensate ageing are currently frustrated by the still limited number of quantitative techniques available for this purpose. The extreme sensitivity to their environment, which confers their biological function, is also what makes protein condensates particularly challenging targets, as it is difficult to prepare them in a state amenable to systematic studies (*17*).

Here, we describe a strategy to overcome this problem and to provide an in-depth characterisation of FUS condensate ageing by considering droplet morphology, material properties and polypeptide conformation as a function of time. We implement this approach through the use of a microfluidic sample deposition method that facilitates the application of surface-based techniques to study biological condensates (*18–20*) (**Figure 1**). We first demonstrate that we maintain the conformational state of condensates in near-physiological conditions on surfaces. Then, using a combination of single-condensate atomic force microscopy (AFM) and bulk infrared spectroscopy (IR), we characterise the heterogeneous changes in condensate structure and mechanical properties over time to reveal local phase transitions in FUS condensates.

**Figure 1.**
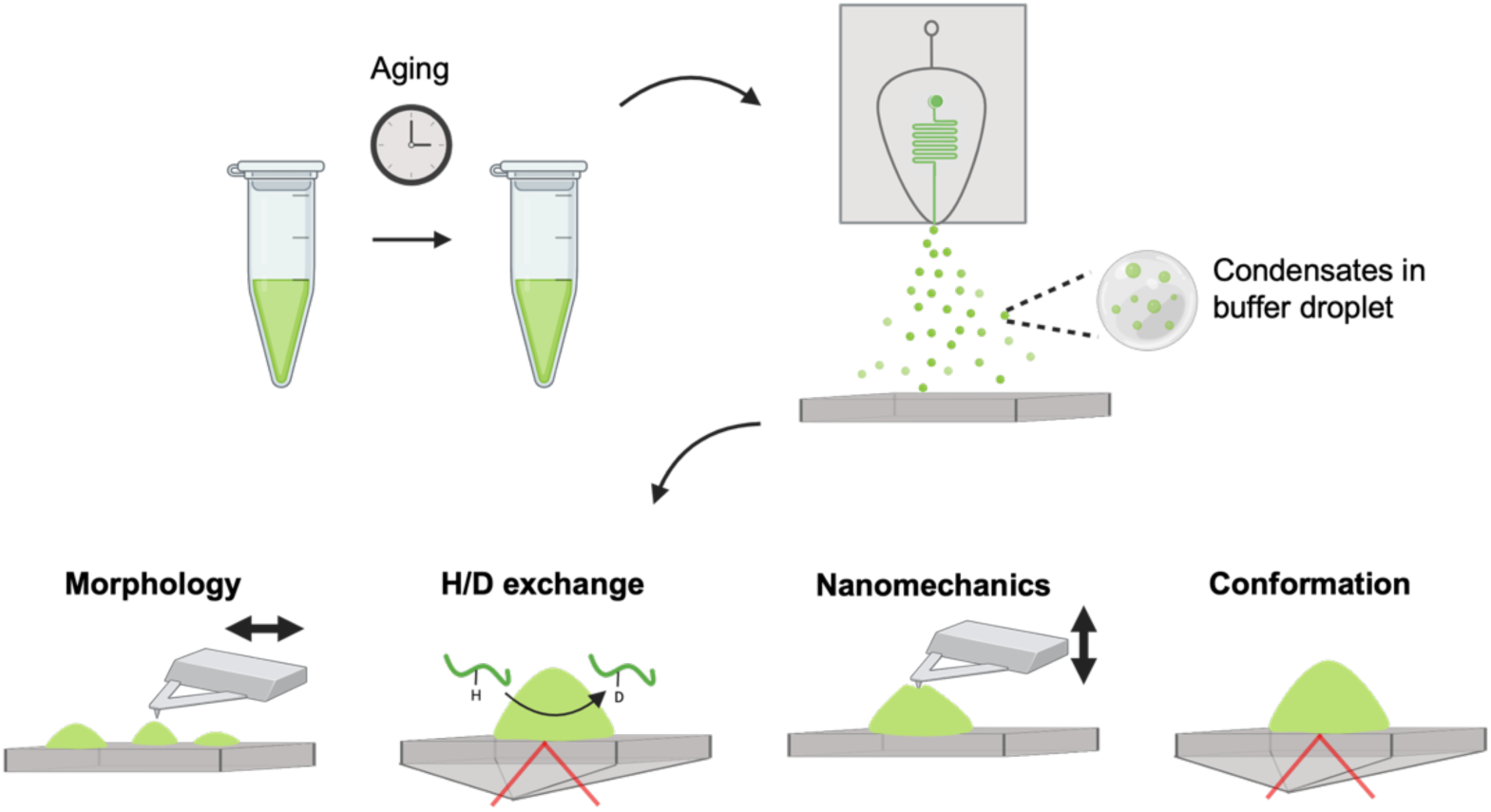
Deposition of ageing FUS condensates via a microfluidic spray device and characterisation of their material properties. Overview of the experiments employed in this study. Condensates were deposited using a microfluidic spray device, which enables their characterisation via surface-based techniques. Four properties of condensates were probed as a function of ageing: (1) 3D morphology via AFM, (2) dynamical properties by hydrogen-deuterium exchange via IR spectroscopy, (3) material properties via AFM nanomechanical mapping, and (4) chemical and structural properties via IR spectrscopy. This combined approach allowed us to generate a detailed model of how phases emerge in space and time within FUS condensates, and how these phases correlate with conformational changes at the protein level.

## Results

### Microfluidic spray deposition of condensates on surfaces

Protein condensates, including those formed by FUS, are notoriously difficult to deposit on surfaces without disrupting the complex interaction network that stabilises the condensed state (*17*). We have previously developed a microfluidic spray deposition method that is capable of preserving the conformational state of biomolecules as in solution (*18*, *20*). Briefly, the method involves the precise generation of picolitre-volume droplets with known dimensions containing the biomolecules, which undergo ultra-fast drying. This ultra-fast drying is on the millisecond timescale, thus minising the time for aberrant conformational changes to occur on the surface (**Figure S1**) (*18*). Therefore, we first sought to assess our ability to deposit the condensates onto surfaces, via microfluidic spray deposition, with preservation of relevant structural features, including morphology and heterogeneity (**Figure 2**). To achieve this, we first imaged condensates in solution via fluorescence microscopy, and compared the same solution deposited via a microfluidic spray device (**Figure S2**). However, under the conditions chosen here, which were selected as a physiological mimic (2 μM FUS, 100 mM KCl) (*21*), the resultant condensates are too small to be readily visualised by light microscopy. Therefore, we turned to high-resolution AFM to robustly assess the morphological properties of the samples.

**Figure 2.**
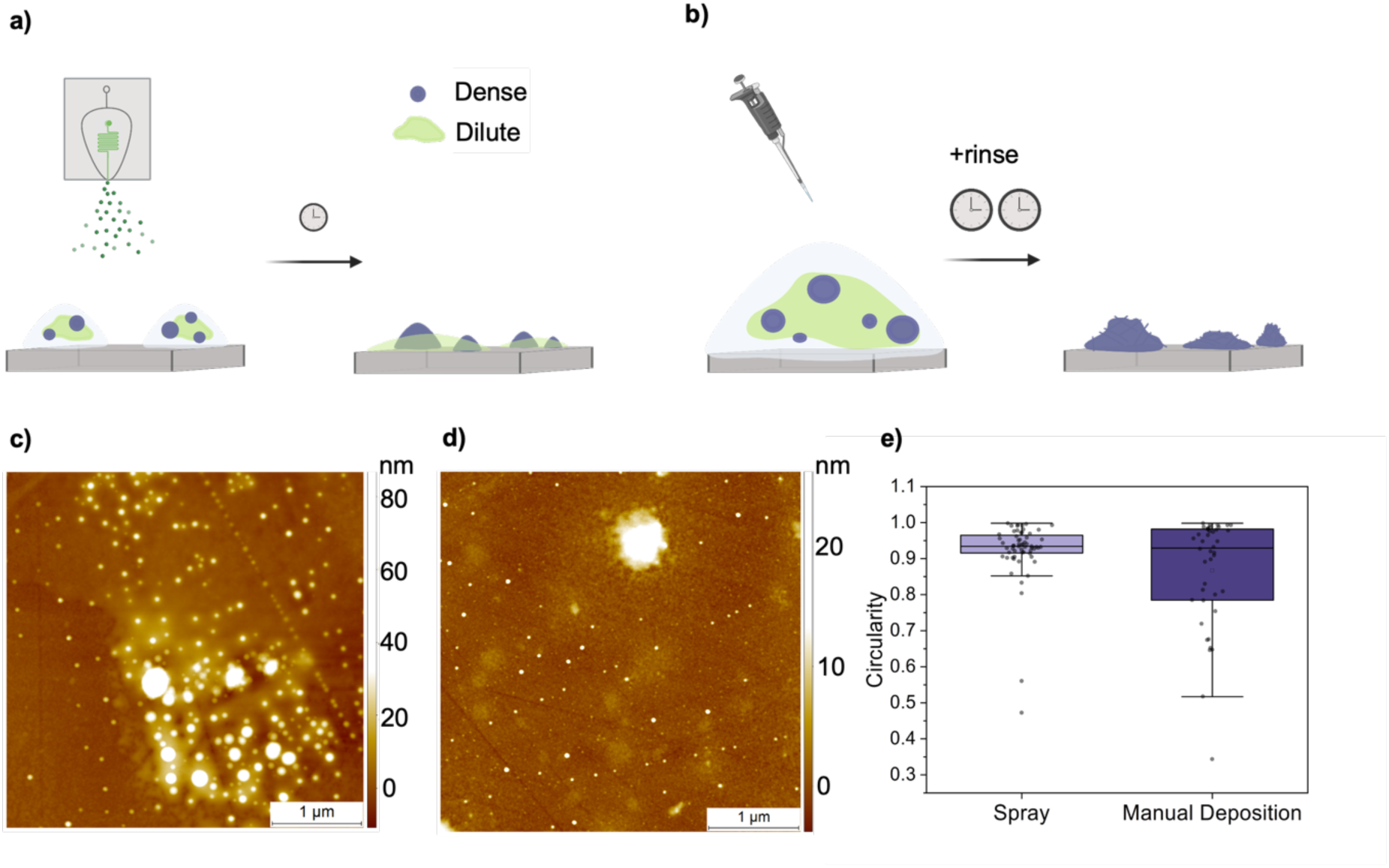
Comparison of surface deposition methods for the characterisation of protein condensates. **(a,b)** Schematics of the sample preparation method for manual deposition (a) and microfluidic spray deposition (b). Manual deposition requires long incubation times (seconds to minutes) and a rinsing step. Spray deposition can be carried out in a single step with short incubation times (tens of milliseconds). **(c,d)** AFM images were acquired for manual (c) and spray (d) deposition, where morphological differences can be observed. **(e)** Circularity was compared for condensates deposited via microfluidic spray deposition and manual deposition.

When we used microfluidic spray deposition, we observed the presence of both the dense and dilute phase, indicating that we are able to preserve the heterogeneity of complex mixtures. We also ensured that the protein was not retained in the microfluidic device by imaging the channels after spraying, where minimal material was observed (**Figure S3**). Further, condensates deposited via microfluidic spray had round morphologies with evidence of fusion events, which are characteristic features of fluid-like condensates in solution. Therefore, we sought to quantify this by measuring the circularity of condensates (**Figure 2e**). We next compared sample properties between microfluidic spray deposition and manual deposition. Briefly, manual deposition involves adding a volume of sample onto a surface, waiting to allow adsorption to the surface, followed by a rinsing step to remove excess material. When deposited manually, in our hands, we no longer observed the presence of the disperse phase, indicating that the rinsing step removes weakly-adsorbed material. Remaining condensates displayed an amorphous, non-circular morphology, which is consistent with surface-induced aggregation of the dense fluid-like phase.

The maintenance of relevant structural features can be rationalised by considering deposition times associated with each method. While manual deposition is typically performed on the second to minute timescale, spray deposition, due to the significant reduction in droplet volume, occurs on the tens of millisecond timescale. To contextualise these considerations, the relaxation time of liquid FUS condensates have been reported to be in the range of tens of milliseconds, which are comparable to the deposition timescales we employ here (*22*). Therefore, there is less time for significant surface-induced structural rearrangement of the protein condensates, which is observed using manual deposition.

Of note, we also spent significant time optimising the imaging surface. As FUS condensates are stabilised by a complex combination of electrostatic and hydrophobic interactions, it is important to identify a suitable surface for imaging. Significant sample deformation was observed for charged surfaces, such as mica, which is typically used in AFM studies (**Figure S4**). In the end, we determined the use of zinc selenide crystals to be ideal, as we did not observe deformations of sample morphology and it has minimal surface defects which can impede high-resolution imaging. However, some alignment of the sample can be observed in surface crevices (**Figure 2c,d**).

### Morphological characterisation of condensate ageing

Having established the ability to maintain relevant structural features of condensates on surfaces, we sought to characterise the changes in morphology as a function of the ageing time (*t_a_*). To this end, FUS samples were incubated, and aliquots were taken at distinct time points (*t_a_* = 0, 2, 4, 8, 24 h) and deposited for surface characterisation (**Figure 1**). We first imaged sample morphology via confocal microscopy, which has the benefit of a large field of view, and thus provides an initial overview of the sample properties (**Figure S2**). At *t_a_* = 0, 2 and 4 h, very few, dim condensates are observed. At *t_a_* = 8 and 24 h, more condensates are observed, which have a higher fluorescence intensity. To characterise this feature in depth beyond the diffraction limit of light microscopy, we also measured condensate morphology via AFM (**Figure 3**).

**Figure 3.**
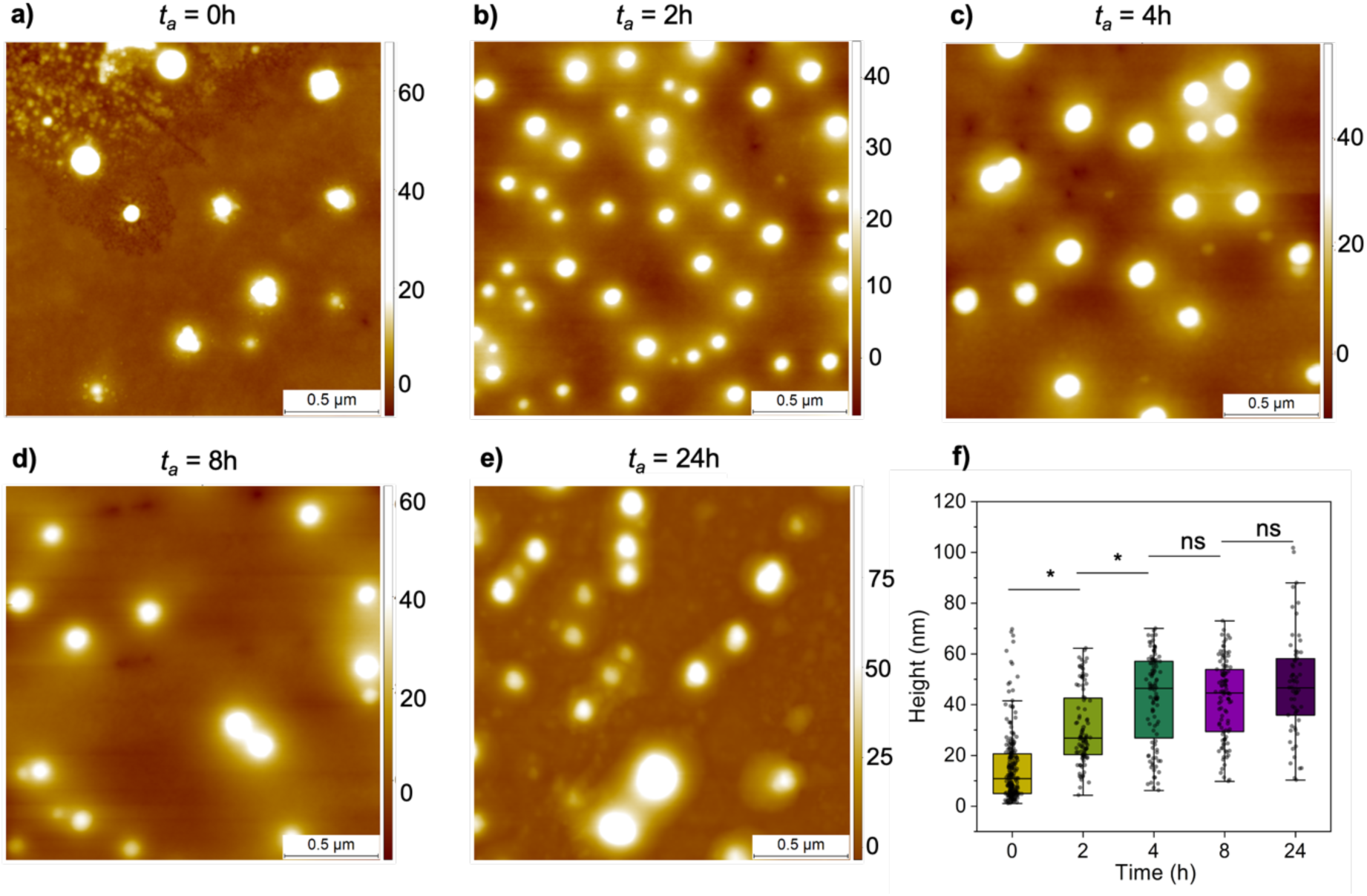
Characterisation of age-related changes in the condensate morphology. **(a-e)** The morphology of condensates was measured via AFM as a function of the ageing time (*t_a_*). Z-scale bar is in nanometres. **(f)** Single condensate statistical analysis was performed on AFM images to determine change in height as a function of ageing time. Mean heights: 0 h 15 ± 14 nm, n = 224; 2 h 31 ± 15 nm, n = 101; 4 h 42 ± 18 nm, n = 104; 8 h 42 ± 15 nm, n = 105; 24 h 48 ± 20 nm, n = 62.

At the initial time point, *t_a_* = 0 h, we observed large condensates as well as numerous small, spherical species, which are consistent with previously described clusters (*23*, *24*). After *t_a_* = 0 h, the size was observed to increase up until *t_a_* = 4 h. After which no significant size increase was further detected, consistent with the loss of condensate growth via fusion, which is a characteristic of fluid assemblies. Therefore, we sought to assess the loss of fluid-like properties via bulk measurements of solidity via the hydrogen-deuterium exchange (HDX) measurements discussed below.

It is also interesting to note that, other than size, there are little obvious changes in condensate morphology as they age. While amyloid fibrils have been reported to form from the low complexity domain of FUS (*25–27*), we have not observed instances of this type of structure in the present work with full-length FUS. This was also confirmed via TEM imaging (**Figure S5**). As FUS solid assemblies have been previously described to reach sizes incompatible with the size of the microfluidic device channels (**Figure S3**), we also manually prepared samples at *t_a_* = 24 h. In these samples, we observe the presence of micrometre-scale fibre-like structures when we prepared aged condensates via manual deposition (**Figure S6**). High-resolution-imaging reveals a loose, mesh-like network in these assemblies, which is not consistent with the formation of highly ordered, cross-β sheet structure typically found in amyloid fibrils (*28*). In our hands, we attribute the formation of these structures to the shear forces from pipetting (*29*, *30*).

### Bulk and single condensate measurements of material properties

To measure the time-dependent loss of dyanamic properties, we turned to HDX via Fourier transform infrared spectroscopy (FTIR), which provides information on the chemical environment, and therefore bonding networks, taking place in the polypeptide backbone (**Figure 4**). This can then be used to infer the fluid-like or solid-like behaviour of condensates. Briefly, hydrogen atoms on the backbone of a dynamic, fluid-like polypeptide will exchange rapidly when the peptide is placed in a heavy water environment (D_2_O), whereas those involved in solid-like structures will be less labile, and therefore the exchange will occur more slowly. Exchange from a hydrogenated state to deuterated state can be monitored via 3 changes in the protein amide peaks: (1) the amide I (∼1650 cm^-1^) peak will experience a shift to lower wavenumbers, which is especially prominent for random coil structure, (2) the amide II (∼1550 cm^-1^) peak will diminish and be replaced by (3) the emergent amide II’ (∼1450 cm^-1^) peak, which increases in intensity with increased deuteration (*31*).

**Figure 4.**
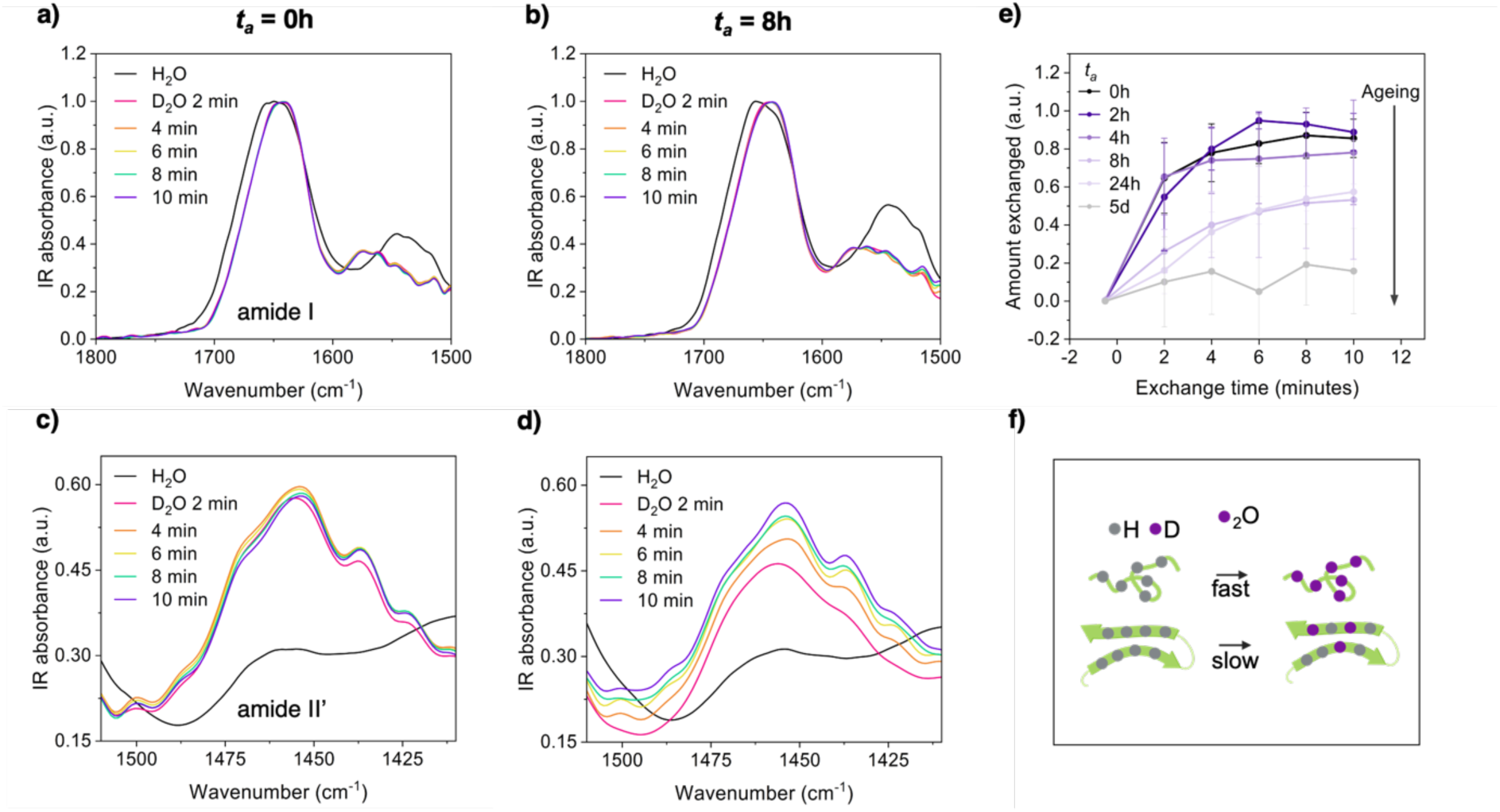
Hydrogen-deuterium exchange provides information on the conformational dynamics within condensates. **(a-d)** Bulk hydrogen lability of protein condensates as a function of *t_a_* was investigated using hydrogen-deuterium exchange via FTIR. Condensates were deposited on the FTIR prism at different *t_a_*, and rate of exchange was measured. HDX can be monitored by changes in the amide I (∼1650 cm^-1^), amide II (∼1550 cm^-1^) (a, b) and amide II’ (∼1450 cm^-1^) (c, d). Black lines represent spectra in H_2_O, and coloured lines show changes in spectra after transfer into D_2_O. Representative spectra are presented for *t_a_* = 0 h (a, c) and 8 h (b, d); the spectra for the other time points can be found in the supplementary information. **(e)** The extent of exchange was monitored over the course of 10 min by taking the integral of the amide II’ peak. **(f)** Schematic displaying the conformation-dependence on backbone proton-lability: protons in random coil regions exchange rapidly, while those involved in hydrogen bonding are less labile (*32*).

Therefore, to probe the progressive loss of fluid-like behaviour of FUS condensates, we quantified the extent of hydrogen-exchange as a function of *t_a_*. At early time points, *t_a_* = 0, 2, and 4 h, backbone hydrogens exchanged rapidly with deuterium in their environment (**Figures 4a,c and S7**). At late time points, *t_a_* = 8, and 24 h, the same shift in the amide I peak was observed (**Figure 4b**). However, exchange occurred to a lesser extent, and more slowly, only sometimes reaching the level of exchange of the early-aged samples after 10 min in D_2_O. This decrease in backbone hydrogen-lability of late-aged condensates may be attributed to polypeptide involvement in hydrogen-bonding networks of a more solid-like assembly, and is consistent with the plateau in condensate size increase observed via AFM.

Therefore, to further understand the emergence of solid-like properties of condensates with ageing, we directly measured the mechanical properties of condensates using AFM. The apparent Young’s modulus, or elastic response, of condensates can be measured by considering how they respond to local deformations by the AFM tip, via the acquisition of force-distance curves (**Figure 5a,b**) (*33*). Elastic response to deformation is a characteristic behaviour of solid materials, therefore the Young’s modulus provides a readout on the emergence of solid-like behaviour. As this method requires a physical deformation of the condensates, we first sought to establish that these measurements were not destructive. To this end, we performed repeat measurements of single condensates to assess for changes in morphology and material properties, both of which serve as an indicator of deformation-induced aggregation. No significant changes in either property were observed (**Figure S8**).

**Figure 5.**
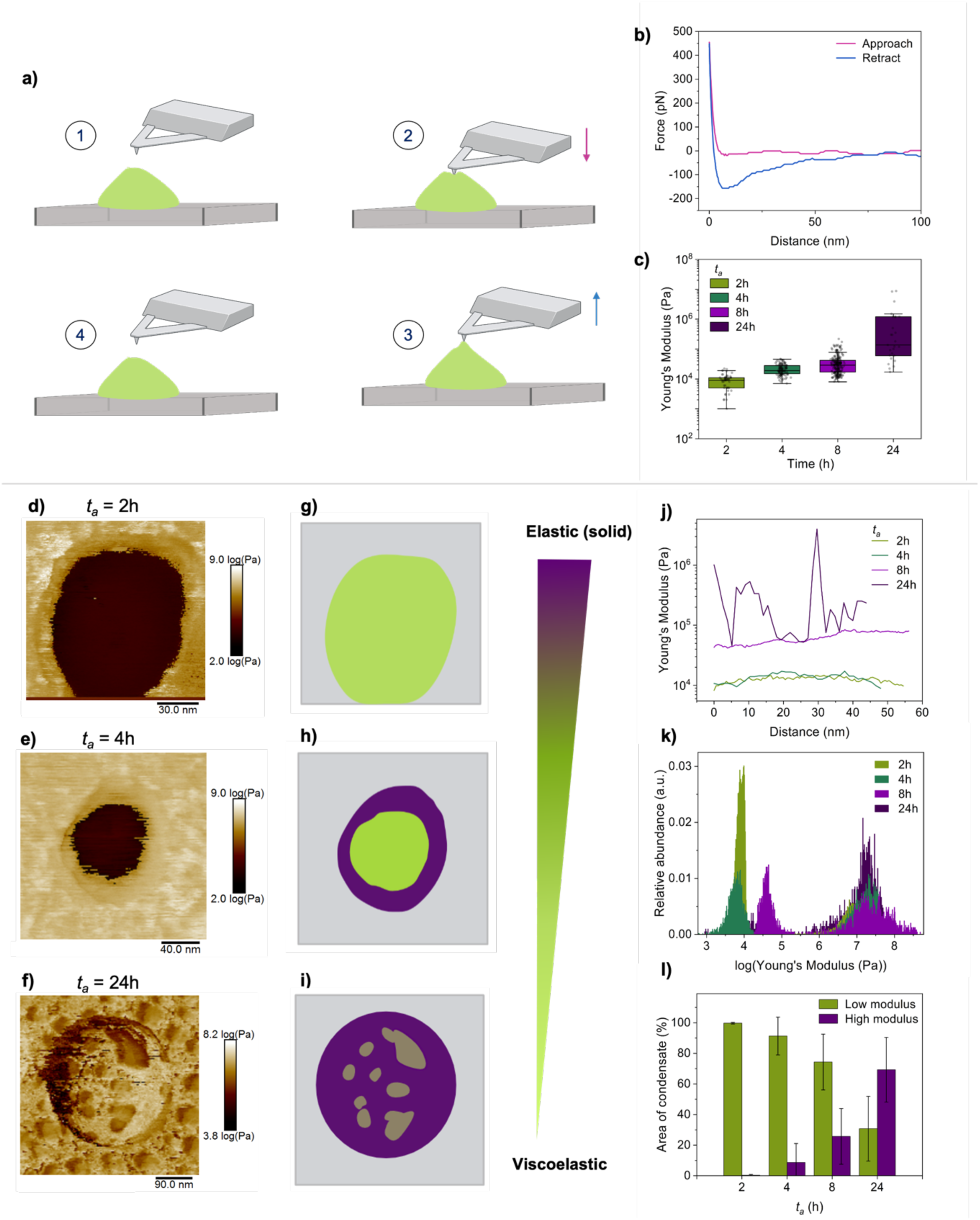
AFM mapping of the material properties of single condensates. **(a)** Schematic showing the procedure of nanomechanical characterisation using an AFM, to generate a force-distance curve. **(b)** A representative force-distance curve is shown for *t_a_* = 24 h. **(c)** Apparent Young’s modulus values of single condensates shown as a function of time. Each point corresponds to the average of three force-distance curves, taken in the centre of condensates (2 h n = 51, 4 h n = 117, 8 h n = 243, 24 h n = 27). **(d-h)** Representative AFM nanomechanical maps of the Young’s modulus in log scale are presented for individual condensates as a function of ageing: *t_a_* = 2 h (d), 4 h (e) and 24 h (f). Corresponding non-quantitative schematics are shown for each nanomechanical map (g-i), to serve as a visual aid. Green represents regions with lower elastic modulus and purple represents regions with higher elastic modulus, a solid-like behaviour. **(j)** Representative cross-sections of the Young’s modulus in the core of condensates are plotted. This reveals a relatively uniform distribution of material properties within the condensate centre up until *t_a_* = 24 h, in which significant heterogeneity emerges. **(k)** A histogram of the Young’s modulus values within a single condensate are shown as a function of ageing. A height threshold was applied to exclude the surface, meaning only the area of the condensate is considered. **(l)** The emergence of the high elastic modulus was quantified as a function of time, by measuring the area of the high modulus phase over the total area of the condensate. Low modulus phase is <1MPa, and the high modulus phase is >1MPa.

Importantly, however, the deposition step being on similar timescales to the relaxation time of very dynamic, liquid FUS condensates, makes them more prone to surface effects (*34*). This, in combination with the lack of elastic response of fluid condensates means that we may not readily report on the mechanical properties of condensates at *t_a_* = 0 h (**Supplementary Note 1 and Figure S9**). Therefore, we report on the elastic modulus as a function of ageing time from *t_a_* = 2 h (**Figure 5c**). These single condensate measurements complement our bulk HDX measurements, in which early time points have elastic modulus values of 9 ± 5 kPa and 21 ± 9 kPa for *t_a_* = 2 h and 4 h, respectively. At *t_a_* = 8 h, the elastic modulus is more variable, with values of 37 ± 31 kPa. At *t_a_* = 24 h, values reach 1.1 ± 2.2 MPa, three orders of magnitude higher than those at *t_a_* = 2 h. These values are consistent with the presence of condensates with mixed material properties, as indicated by the large range of measured elastic values. These values are also in a similar range to previously described solution-based measurements, albeit higher, which may be explained by the presence of the surface (*12*).

Therefore, we sought to understand the heterogeneity of the material properties within single condensates via nanomechanical mapping, which provides nanometre spatially-resolved mechanical information (**Figure 5d-f, S10**) (*35*). At *t_a_* = 2 h, we observed condensates with uniform, low elastic modulus mechanical properties. At *t_a_* = 4 and 8 h, we began to observe the emergence of a high elastic modulus, solid-like shell at the edge of the condensates. Importantly, AFM nanomechanical measurements display an inherent artefact of edge effects, in which mechanical properties of a sample are not accurately measured at boundaries (*36*). This is particularly relevant in instances of soft samples on hard supports, as used here, as the measured elastic moduli will include a combination of contributions from the soft sample and the hard substrate below, due to the round shape of condensates leading to lower heights at their edges. Therefore, to decouple edge effects from the phenomena of interest, we measured the thickness of the high-modulus shell as a function of time. One would expect the condensates at *t_a_* = 2 h to display the largest edge effects, due to their low elastic moduli. However, in fact we observed the opposite, with a thicker shell at *t_a_* = 4 and 8 h (**Figure S11**). Therefore, even while considering edge effects, there is a high-modulus phase which forms at the condensate-solvent interface as condensates age, which is consistent with previous observations (*12*, *37*). Finally, at *t_a_* = 24 h, we observed significant heterogeneity across the whole condensate, with co-existing low and high-modulus phases (**Figure 5f,j**).

Having established the spatially-inhomogeneous distribution of low and high-elastic modulus phases within single condensates, we sought to quantify the emergence of the high-elastic modulus phase as a function of time. Therefore, we plotted the distribution of Young’s modulus values across whole condensates (**Figures 5k and S12**) At *t_a_* = 2 h, there is one dominant peak corresponding with a low elastic modulus phase. At *t_a_* = 4 h, there are two dominant peaks, indicating the presence of a dominant, low elastic modulus phase in the condensate core, and a less abundant, high elastic modulus phase at the condensate-solvent interface.

At *t_a_* = 8 h, this same high elastic modulus phase exists. However, the elastic response of the soft core increases. Finally, at *t_a_* = 24 h, the dominant peak is the high elastic modulus phase. These results indicate that a high-elastic modulus shell formation precedes a global increase in the elastic response of the centre of the condensates. This high-modulus phase then propagates heterogeneously through the condensate core.

Finally, the identification of well-separated peaks corresponding to distinct phases allowed us to quantify the emergence of the high-modulus phase as a function of ageing time (**Figure 5l**), providing information on its kinetics of formation. This also allowed us to understand precisely what phase is present at each time point, enabling for robust correlation between material properties and polypeptide behaviour measured via bulk techniques. This is important for identifying the structural and conformational features of particular physical states, which we establish in the following section.

### Age dependent changes in conformation and secondary structure

Having characterised changes in morphology and material properties of FUS condensates in physiologically-relevant conditions, we next sought to determine whether there were corresponding changes in secondary structure and conformation via FTIR. The use of this technique is greatly facilitated by the microfluidic spray deposition. Ultra-fast deposition times minimise salt crystallisation, thereby reducing the contribution of buffer salts to the spectra (*18*). This affords a significant increase in spectroscopic sensitivity, such that we can extract relevant features which would typically be difficult to distinguish from noise, and measure secondary structure in relevant high salt conditions (>50 mM NaCl). This is important, as FUS phase behaviour is intrinsically linked to the protein concentration and ionic strength (*38*).

At *t_a_* = 0, 2, and 4 h, the spectrum is dominated by random coil/α-helical structure, which has a characteristic broad amide I peak at 1657 cm^-1^ (**Figure 6a,c,d**). This is also confirmed by a downward shift of ∼10 cm^-1^ in D_2_O, which is characteristic of random coil structure (**Figure 3a**). These structural features are in line with solution-structures of FUS, which reveal a long, intrinsically disordered domain as well as a α-helical, RNA-recognition domain (*39*). This is also consistent with reports that the transition from the dilute to dense phase is not associated with any measurable change in secondary structure (*40*). Additionally, the width of the amide I peak can be used to understand conformational flexibility. As each spectrum represents the averaging of snapshots of the conformations of each polypeptide in bulk, a broad peak indicates the averaging of many conformations. Spectra of condensates at these earlier time points of ageing all display broad peaks, indicating a high degree of conformational flexibility, which is expected of a fluid-like state (**Figure 6b**) (*8*). Of note, there is a native β-sheet peak at 1635 cm^-1^, which we assigned to the β-barrel of GFP tag (*41*). A tagged protein was used to enhance protein solubility for ease of handling and to facilitate comparison with light microscopy experiments.

**Figure 6.**
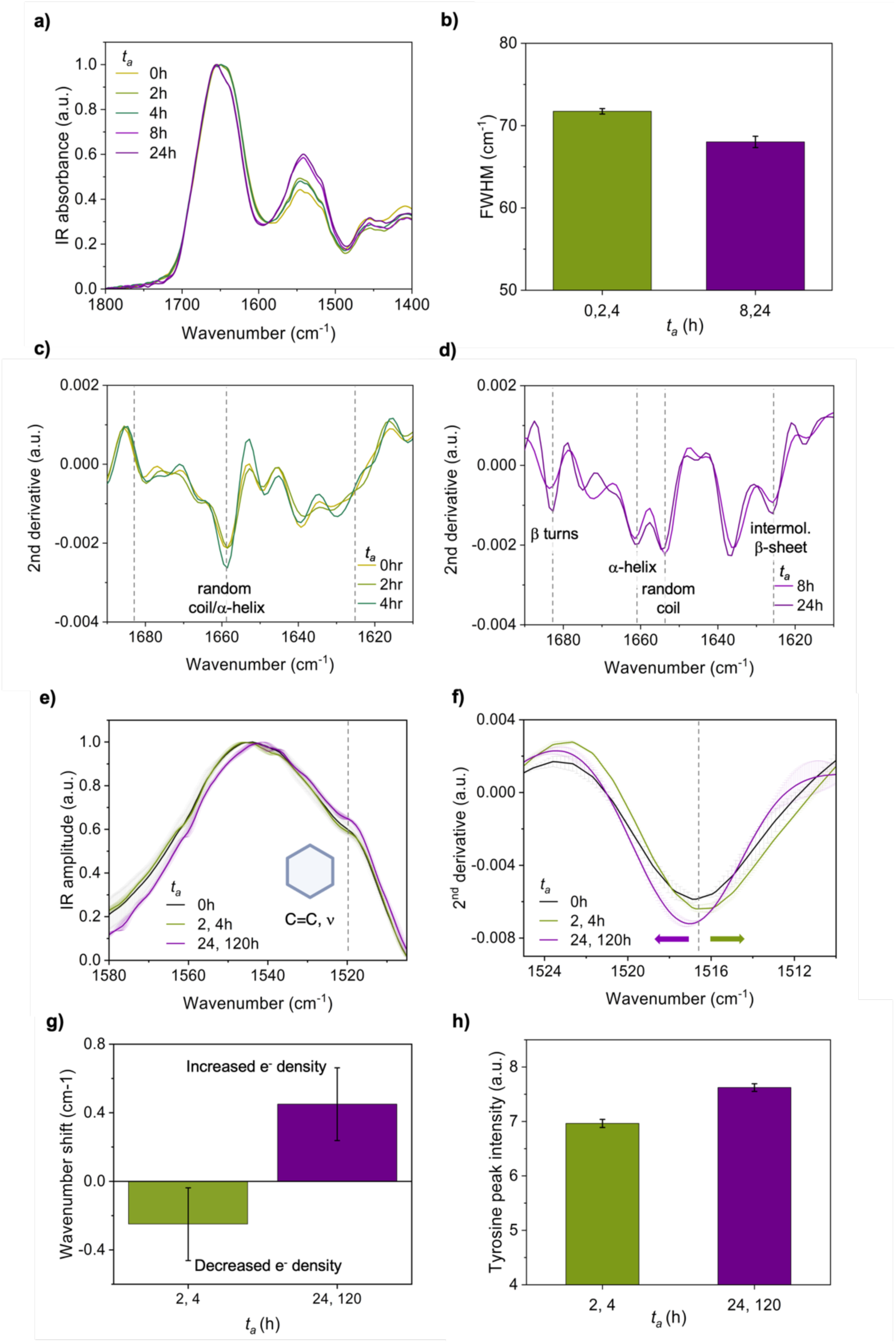
Secondary structure changes associated with the gel to solid transition of FUS. **(a)** The secondary structure of FUS condensates were measured as a function of time using FTIR, at *t_a_* = 0 to 24h. **(b)** The width of the amide I peak was measured, FWHM = full width at half maximum. **(c-d)** Th second derivative of the amide I bands are presented for early, *t_a_* = 0, 2 and 4 h (c) and late, *t_a_* = 8 and 24 h (d) condensates. Differences in secondary structure components are highlighted based on characteristic absorption peaks. **(e)** Tyrosine residues, and their chemical environment, can be measured via FTIR. The tyrosine absorption peak for the liquid/dense phase is at ∼1517 cm^-1^, indicated by the black dashed line. Tyrosine peaks were also recorded for *t_a_* = 2 and 4 h (green) and solid *t_a_* =24 h and 5 d (green) samples. A different chemical environment will result in shifts of the peak. **(f)** The second derivative of the tyrosine peak reveals downward and upward shifts for early-aged condensates (*t_a_* = 2 and 4 h, green) and late-aged condensates (*t_a_* = 24 and 120 h, purple), respectively, relative to the peak position of at *t_a_* = 0 h (black). The black dashed line indicates the peak position at *t_a_* = 0 h. **(g)** The shifts in the tyrosine peak relative to *t_a_* = 0 h were quantified. **(h)** The intensity of the tyrosine peak was measured by considering the integral of the peak. Differences in the peak intensity at *t_a_* = 24 and 120 h indicates a different chemical environment.

As *t_a_* increases, corresponding with the formation of a high elastic modulus phase, we observed new features emerging in the spectra. Firstly, we observed two peaks emerge at 1625 cm^-1^ and 1682 cm^-1^, indicating the presence of intermolecular β-sheet and β-turns, respectively (**Figure 6c**). The intensity of both these peaks increases between *t_a_* = 8 and 24 h. Also, where we initially saw one major peak which represented a combined random coil/α- helical structure, we now observe two individual peaks at 1660 cm^-1^ (α-helix) and 1653 cm^-1^ (random coil), the assignment of which is assisted by comparing spectra in H_2_O and D_2_O (**Figure S13**). We attribute this peak divergence to a decrease in the amount of random coil structure, which is replaced by β-sheet structure. We can understand the nature of the β-sheet assemblies here by considering both the peak position and width, which provides information on chemical environment and conformational heterogeneity. Peaks of intermolecular β-sheets are typically in the range of 1625-1610 cm^-1^, with higher density of intermolecular hydrogen-bonding resulting in a shift to lower wavenumbers (*32*). Furthermore, the peak width only decreased by ∼1.5%, compared to spectra from *t_a_* = 0, 2 and 4 h, which is within the error of the measurements. Taken together, this is consistent with a high degree of conformational heterogeneity being retained, even in the solid-phase.

Having established the presence of heterogenous β-sheet assemblies, we next sought to assess the interaction network involved in the solid-like phase. FTIR spectroscopy is also capable of measuring the chemical environment of side chains, and therefore can be used to understand their interactions. Here, we focussed on tyrosine residues, which are enriched in the LTD, and are believed to be key drivers of phase separation due to their involvement in cation-π interactions with arginine residues (*9*, *10*). Tyrosine can be monitored via the C=C stretching vibration of the aromatic ring, which has a characteristic peak at ∼1517 cm^-1^ (**Figure 6e**) (*42*). To increase our certainty in the peak assignment, we also compared spectra to those acquired in D_2_O, where we see the expected downward shift to ∼1514 cm^-1^ (**Figure S14**) (*42*). At *t_a_* = 2 and 4 h, we observed a downward shift relative to the peak at *t_a_* = 0 h, which is consistent with a decrease in the electron density in the aromatic ring (**Figure 6f,g**) (*42*). We attribute this to an increase in the electron donation from the tyrosine to an electron accepting group, which may be consistent with an increase in the density of cation-π interactions (*43*). At later time points of ageing, corresponding with the formation of solid-like, β-sheet assemblies, we see the opposite trend, with an upward shift of the tyrosine C=C peak, suggesting an increase in the electron density in the aromatic ring (**Figure 6f,g**). At *t_a_* = 8 h, in which both dynamic and solid phases are observed (**Figures 5 and S12**), the peak position is approximately the same as at *t_a_* = 0 h, with only a slight downward shift, likely indicating an averaging of the two relevant interactions in the co-existing dynamic and solid phases observed in the condensates (**Figure S15**). We also observe an increase in the intensity of the tyrosine peak at later time points of ageing (**Figure 6h**), again suggesting a different chemical environment of these residues. We therefore suggest a different form of interaction appears to be relevant in stabilising the fluid and solid phases.

## Discussion

We have described the results of a study of the phase behaviour of full-length FUS condensates upon ageing, which was carried out using a microfluidic spray deposition method that enables the accurate characterisation of the condensates on surfaces. We have demonstrated that it is possible to preserve relevant conformational features of FUS condensates on surfaces, thus enabling us to perform high-resolution structural and mechanical characterisation of the ageing behaviour of the condensates in physiologically-relevant conditions. Furthermore, by performing structural studies using surface-based techniques, we are no longer limited by polypeptide length, as some solution-based methods are, allowing us to study full-length FUS. This experimental procedure made it possible to build on the valuable insights provided from structural studies on individual domains, such as the low complexity domain, giving us an understanding of how the overall properties of the full protein sequence may alter the ageing behaviour of FUS condensates (*44*).

The enhanced capability of studying condensates on surfaces has enabled us to generate a detailed model of how solid-like features temporally and spatially emerge within initially liquid-like condensates, and how these phase changes correlate with conformational changes at the molecular level (**Figure 7**). Initially, we have observed the presence of uniform condensates with fluid-like properties imparted by the intrinsically-disordered conformation of component protein molecules. An increasingly elastic, solid-like, intermolecular β-sheet-rich phase then forms at the condensate-solvent interface, while the condensate core remains fluid-like, which is consistent with previous reports (*12*, *13*). Subsequently, there is also an increase in the density of intermolecular interactions between intrinsically-disordered FUS molecules within the fluid core, resulting in an increasing elastic response. Finally, the solid phase heterogeneously permeates from the condensate edges through the aged fluid core. Spectral analysis has revealed conformational heterogeneity and low-density intermolecular interactions between β-sheets within the solid phase, suggesting that the solid structures formed here are structurally distinct from amyloid fibrils.

**Figure 7.**
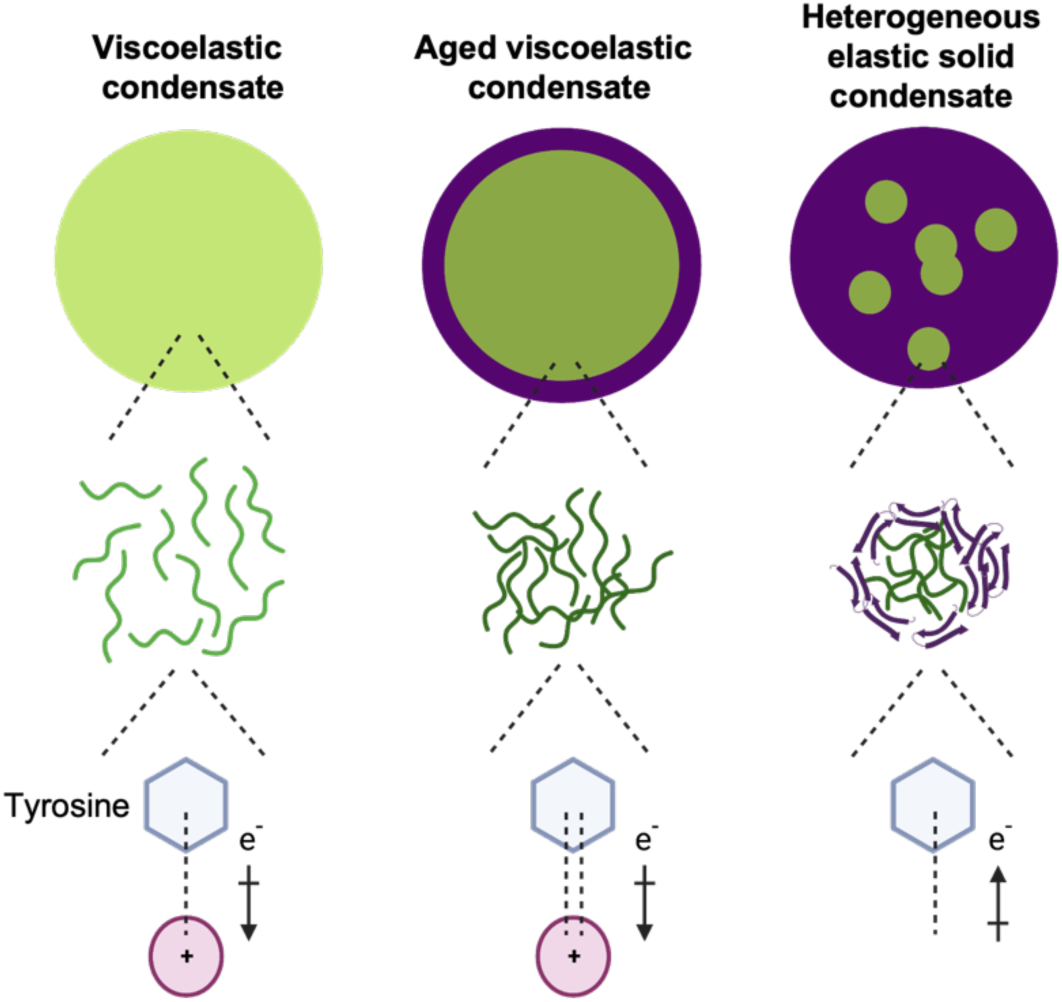
Local phase transitions in ageing FUS condensates. FUS condensates can be initially characterised as viscoelastic materials, as they exhibit both viscous (i.e. liquid-like) and elastic (i.e. solid-like) characteristics when undergoing deformation (*34*). Upon ageing, the liquid-like behaviour is gradually lost and more solid-like characteristics emerge. Initially uniform condensates are composed of intrinsically-disordered polypetides. An increasingly elastic, solid-like, β-sheet-rich phase then forms at the condensate-solvent interface, while the condensate core remains disordered. Subsequently, there is also an increase in the density of intermolecular cation-π interactions between intrinsically-disordered FUS molecules within the core, resulting in an increasing elastic response. Finally, the amorphous solid phase heterogeneously permeates from the condensate edges through the core.

We have also observed that tyrosine residues adopt a different conformation in these solid assemblies, indicating that they are involved in different interaction networks. We note that tyrosine residues have been shown to be required for transcriptional activity, and thus their different molecular environment in solid assemblies may affect their accessibility, perhaps in connection with the pathological nature of aged FUS assemblies (*11*, *45*).

The challenges in obtaining detailed characterisations of the physical changes in ageing condensates have so far hindered our understanding of how phase behaviour plays a role in biological function. Of particular interest is an intermediate, broadly-defined gel state which has been described frequently in the literature for a variety of biomolecular condensates. Such assemblies typically show decreased protein mobility within condensates, identified by fluorescence recovery after photobleaching (FRAP), or reduced fusion rates (*46*). Many functional assemblies possess this mobility behaviour, highlighting the relevance of complex material properties in defining condensate form and function (*47–51*). We observe that this behaviour reported in the literature would be well described by the aged fluid condensate architecture that we have described here - in particular, condensates possessing a solid-like shell and an aged fluid-like core (*52*). We have correlated the increase in density of intermolecular interactions between intrinsically disordered proteins with changes in the bulk elastic response of condensates, thus presenting a physical basis to explain the properties of this gel-like material state. Thus, we expect that an understanding of the heterogeneous phase distribution within condensates will enable more accurate definitions of these biologically-relevant states.

Great efforts are being devoted to the identification of how the physical properties of condensates are linked with their biological functions, as well as their aberrant phase transitions in disease (*16*, *53*). A detailed knowledge of the physical and chemical properties of biological condensates creates a framework to understand how modulators, either biological or chemical, may alter phase transitions. Thus, we hope that this work will serve as a platform to clarify in greated detail the role of phase behaviour in cellular function and dysfunction. Finally, and more generally, we anticipate that the ability to characterise biological condensates on surfaces will create many opportunities to better understand the properties and behaviours of these complex assemblies.

## Materials and Methods

### Production and purification of full-length FUS protein

FUS protein was purified as previously described (*10*). Constructs encoding FUS residues 1-526 were cloned into pACEBac2 vector with an EmGFP C-terminal tag. Proteins were expressed and purified from insect Sf9 cells infected with the baculovirus. After four days of infection cells were harvested by spinning at 4000rpm for 30 minutes. Cell pellets were mixed with the resuspension buffer containing 50 mM Tris, 1 M KCl, 0.1% CHAPS, 1 mM DTT, 5% glycerol at pH 7.4, and proteins purified using three steps purification scheme including, Ni-NTA affinity column, Amylose affinity column followed by size exclusion chromatography in the buffer containing 50 mM Tris, 1 M KCl, 1 mM DTT, 5% glycerol at pH 7.4.

### Fabrication of microfluidic devices

A two-step photolithographic process was used to fabricate the master used for casting microfluidic spray devices(*54*). In brief, a 25 μm thick structure was fabricated (3025, MicroChem) was spin-coated onto a silicon wafer. This was then soft-baked for 15 min at 95 °C. An appropriate mask was placed onto the wafer, exposed under ultraviolet light to induce polymerization, and then post-baked at 95 °C. A second 50 μm thick layer (SU-8 3050, MicroChem) was then spin-coated onto the wafer and soft-baked for 15 min at 95 °C. A second mask was then aligned with respect to the structures formed from the first mask, and the same procedure was followed, i.e. exposure to UV light and post-baking for 15 min at 95 °C. Finally, the master was developed in propylene glycol methyl ether acetate (Sigma-Aldrich) to remove any photoresist which had not cross-linked. A 1:10 ratio of PDMS curing agent to elastomer (SYLGARD 184, Dow Corning, Midland, MI) was used to fabricate microfluidic devices. The mixture was cured for 3 h at 65 °C. The hardened PDMS was cut and peeled off the master. The two complementary PDMS chips are then activated with O2 plasma (Diener Electronic, Ebhausen, Germany) and put in contact with each other and aligned precisely such that the gas inlet intersects with the liquid inlet to form a 3D nozzle (*18–20*).

### FUS sample handling

Aliquots of FUS solution was thawed immediately before use. Phase separation was induced in an Eppendorf tube by lowering the salt concentration, via sample dilution to a final concentration of 100 mM KCl, 25 mM Tris. Aliquots were then taken from the solution at desired time points (i.e. 0, 2, 4, 8 and 24 h) and promptly deposited via microfluidic spray deposition (described below). For measurements with manual sample deposition, 10 μl of sample was deposited on the surface and allowed to adsorb for 5 min. The sample was then rinsed with MilliQ water and dried with a gentle stream of nitrogen gas. For all measurements on the 5-day aged sample, manual deposition was performed, due to the viscosity of the sample preventing it from passing through the device channels.

### Microfluidic spray deposition

Prior to introduction of sample, each device was tested and washed with MilliQ water for 5 min. Sample was then loaded into 200 μL air-tight glass syringes (Hamilton) and driven into the spray device using a syringe pump (Harvard apparatus). Solutions containing sample were pumped into the device with a maximum flow rate of 100 μl/h to minimise sample shearing, while the nitrogen gas inlet pressure was maintained at 3 bar. Deposition was conducted for a maximum of 10 s at a distance of 3.5 cm to ensure coalescence of droplets did not occur. Samples were sprayed directly onto the relevant surfaces (i.e. ZnSe crystals, FTIR prism) with no further washing steps required before measurements.

### Confocal microscopy

FUS condenates were deposited via microfluidic spray deposition, or via manual pipetting of ∼20 μl onto a glass coverslip. Images were taken using a confocal microscope (Leica, TCS SP8). Excitation/Emission wavelengths of 405nm/510nm were used respectively to image the condensates. All images were taken under ambient conditions, and acquisition was initiated immediately after deposition, in order to minimise any evaporation, in the case of manual deposition.

### Atomic force microscopy

Samples were deposited onto ZnSe crystals via microfluidic spray deposition. High-resolution morphology measurements were performed in ambient conditions on an NX10 AFM operating in non-contact mode (Park Systems, South Korea). PPP-NHCR probes were used, with a spring constant of 42 N/m and with a nominal tip radius of ∼10 nm. Images were acquired at a scan rate of 0.3-0.5 Hz to minimise forces applied to the delicate condensates. Nanomechanical characterisation was performed in 25 mM Tris buffer using a MultiMode 8 (Bruker, USA) operating in either force-volume or quantitative nanomechanical mapping mode. ScanAsyst Fluid probes were used, with a spring constant of 0.7 N/m and a nominal radius of ∼20 nm. Images were acquired at scan rates of 0.3-0.5 Hz, at 512×512 pixels. Probes were calibrated using the thermal tune method. Peak force values were chosen between 200-1000 pN. Deformation was monitored to ensure consistent indentation values across condensates with different material properties, and to ensure indentation depths did not result in excessive sampling of the substrate. Only condensates above 30 nm in height were considered. All measurements were performed at room temperature. Scanning Probe Image Processor (SPIP) (version 6.7.3, Image Metrology, Denmark) software was used for image flattening and single condensate statistical analysis. Force-distance curves were analysed in Nanoscope Analysis software (Bruker, USA). A Hertz-DMT contact mechanic model was used to fit the contact region of withdrawal curves. The contact region of the withdrawal curve was analysed via second derivative analysis; those curves which showed a biphasic response were rejected, as this was taken as an indication of contribution from the hard substrate.

### FTIR and HDX

Measurements were performed on a Vertex 70 FTIR spectrometer (Bruker, USA) equipped with a DiamondATR unit and a deuterated lanthanum a-alanine-doped triglycine sulfate (DLaTGS) detector. Each spectrum was acquired with a scanner velocity of 20 kHz over 4000 to 400 cm-1 as an average of 256 scans. Thin films were deposited on the prism using microfluidic spray deposition. For traditional sample deposition methods, 10 μl of sample was deposited to create a thin protein film covering the prism. The sample was allowed to adsorb for 5 min, then subsequently blotted and rinsed with 5 μl MilliQ water. New background spectra were acquired before each measurement. All spectra were normalised and analysed using OriginPro (Origin Labs). To determine the secondary structure composition of proteins, a second derivative analysis was performed. Spectra were first smoothed by applying a Savitzky-Golay filter. For HDX experiments, the samples were prepared using microfluidic spray deposition. The prism, with deposited protein thin film, was first flushed with H_2_O vapour to serve as a baseline, as sample film swelling can affect acquired spectra. Vapour was produced by bubbling N_2_ gas through H_2_O to a home-built chamber around the prism. The input lines to the chamber were then switched to flush the prism with D_2_O vapour and measurement started immediately. The integral of the amide II’ peak intensity (1450 cm^-1^) was used to monitor exchange over time. The presence of residual H_2_O in the chamber was monitored via the peak at ∼3300 cm^-1^ to remove the confounding variable of differing D_2_O amounts.

## Acknowledgements

We gratefully acknowledge Dr. Heather Greer for assistance with TEM, supported by EPSRC Underpinning Multi-User Equipment Call (EP/P030467/1). We also thank Dr. Michael Metrick for helpful advice regarding FTIR hydrogen-deuterium exchange experiments, and Dr. Selen Manioglu and Prof. Daniel Müller for helpful advice regarding multi-parametric nanomechanical mapping experiments.

