## Supplementary Information for "Nanoscale profiling of evolving intermolecular interactions in ageing FUS condensates"

### Supplementary note 1: Nanomechanical properties of non-aged condensates

Despite evidence suggesting that we are able to maintain secondary structure of protein condensates, due to the drying step being on similar timescales to the relaxation time of liquid FUS condensates (*1*), it may not be fully justified to report on the mechanical properties of non-aged, highly dynamic condensates ($t_a$ = 0 h). This is confirmed by direct nanomechanical measurements of condensates at $t_a$ = 0 h, for which Young's modulus values are higher than that expected for a liquid-like assembly (**Figure S10**). Further, attempts to dissolve condensates at $t_a$ = 0 h after spray deposition using hexane-diol did not result in complete dissolution, indicating that we do not maintain the liquid-like mechanical properties of the assemblies (**Figure S10**). However, the timescales of drying and deposition are at least three orders of magnitude faster than the relaxation time for aged FUS condensates (tens of ms vs ~4.8 s), thus enabling us to accurately report on the changes in mechanical properties as a function of time from $t_a$ = 2 h (*1*).

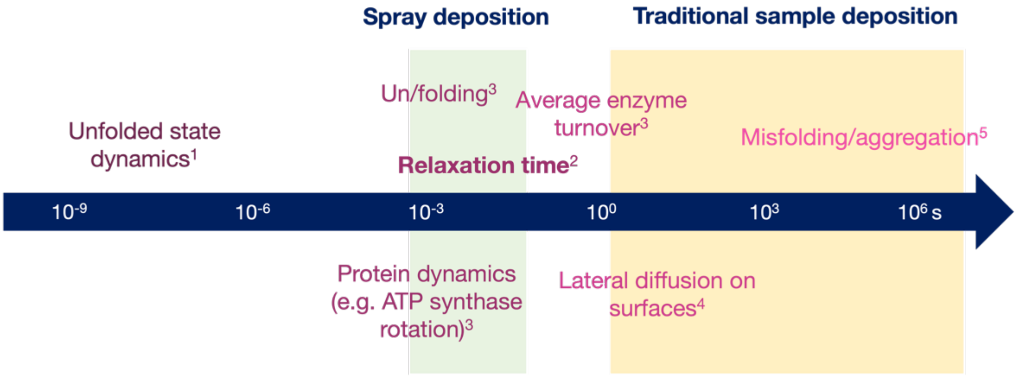


**Figure S1. Timescales of protein dynamics.** The timescales of drying in spray deposition versus manual deposition are contextualised by considering timescales of protein dynamics. Spray deposition times are comparable to reported relaxation times of biomolecular condensates. Values taken from the following references: (*2*)[1], (*1*)[2], (*3*)[3], (*4*)[4], and (*5*)[5].

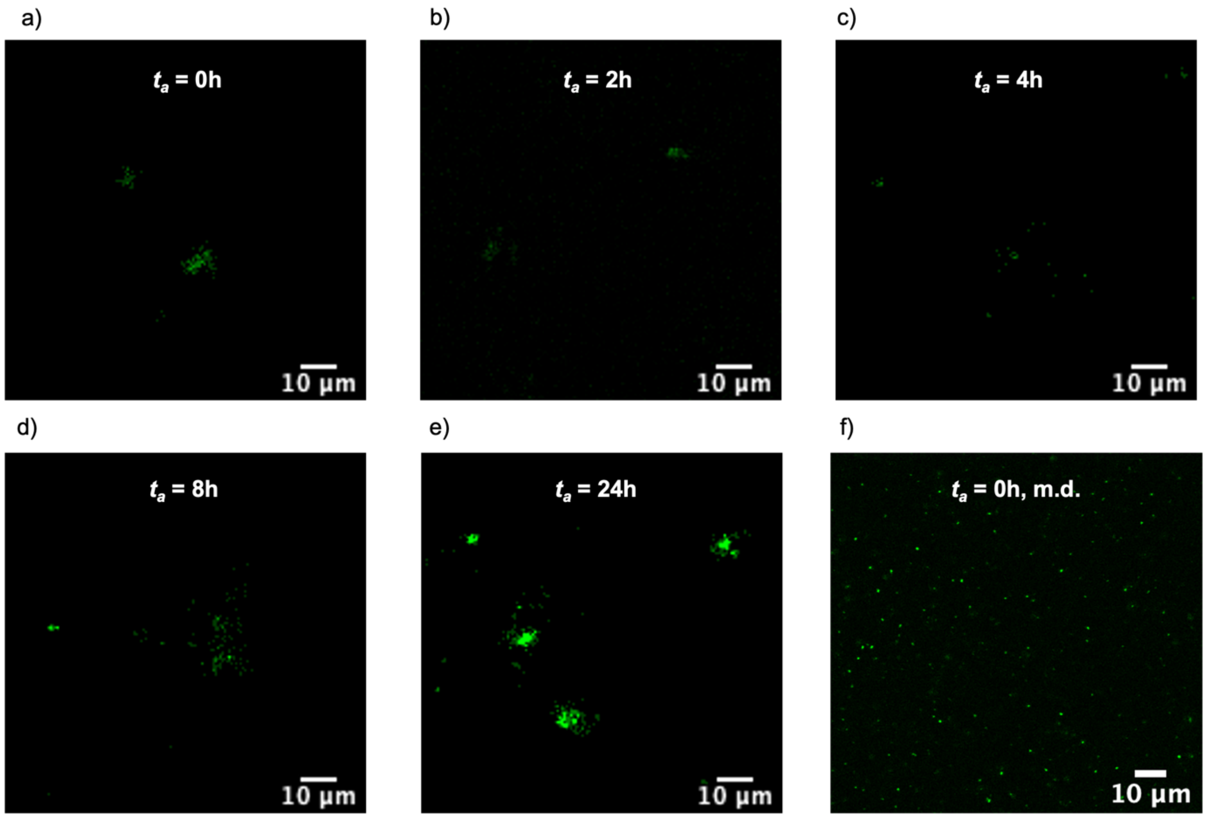


**Figure S2. Confocal images of FUS condensates at various ageing times. (a-e)** Condensates deposited via microfluidic spray were imaged at $t_a$ = 0 (a), 2 (b), 4 (c), 8 (d), and 24 h (e). Very few, faint condensates are observed at early ageing times ($t_a$ = 0, 2 and 4 h), and more, brighter condensates are observed at late ageing times ($t_a$ = 8 and 24 h). **(f)** Samples were also imaged in solution, deposited via pipetting.

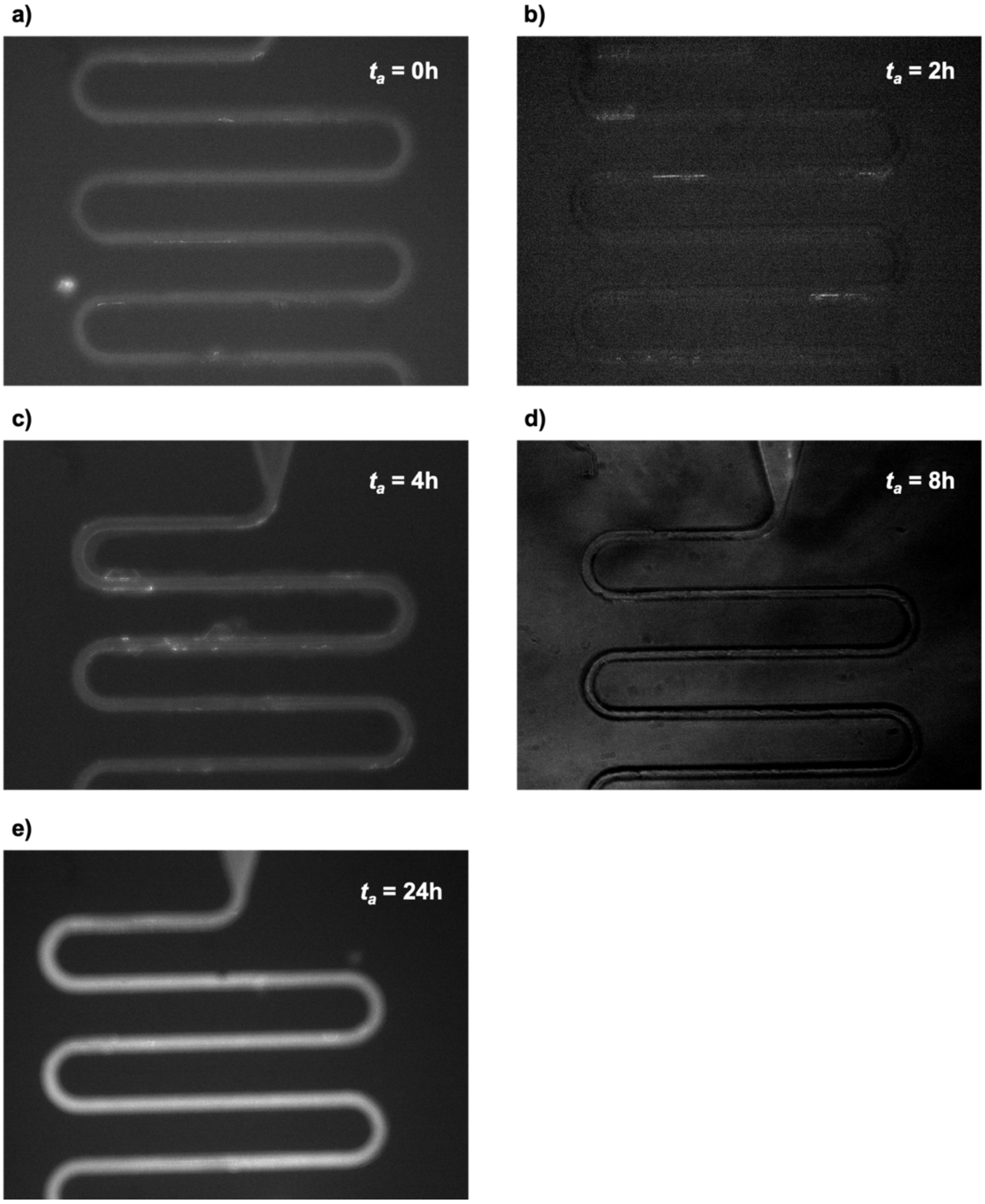


**Figure S3. Fluorescent images of FUS deposition within microfluidic device channels. (a-e)** Channels of the microfluidic spray devices were imaged after sample deposition to assess the extent of bio-fouling. Low levels of fluorescence intensity at early time points, $t_a$ = 0 h (a), and 2 h (b) indicate little material is lost in the device. At a late time-point, $t_a$ = 24 h (e,f), there is higher fluorescence intensity, indicating that some material is retained in the channels (e).

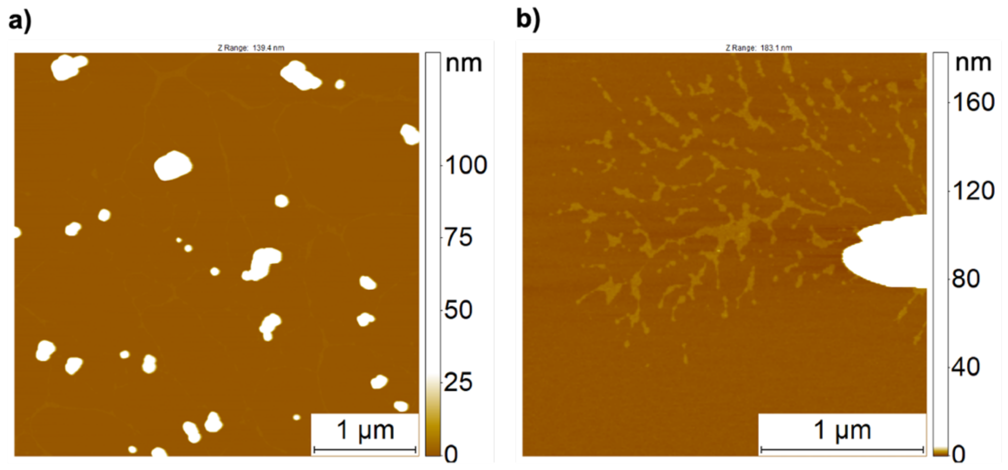


**Figure S4. AFM imaging of deformation of FUS condensates upon deposition on mica surfaces. (a-b)** Sample deformation was observed when FUS condensates were deposited on charged mica substrate. Condensates appeared to have amorphous conformations (a) and appeared to wet the surface (b).

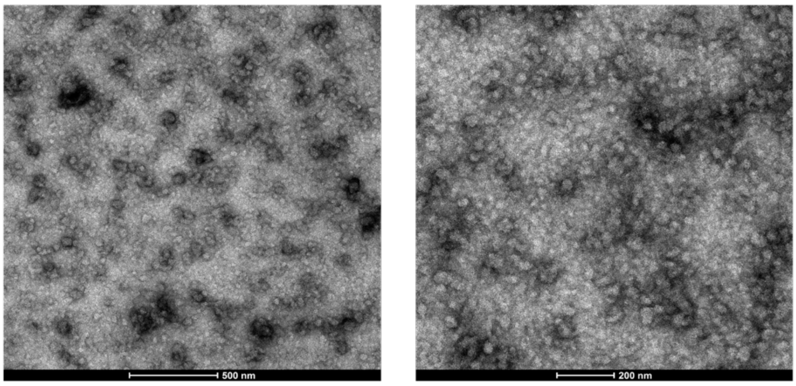


**Figure S5. TEM images of solid-like aged FUS condensates.** Condensates were deposited and imaged via TEM at $t_a$ = 24 h. Condensates had an amorphous appearance, and no amyloid fibrils were observed.

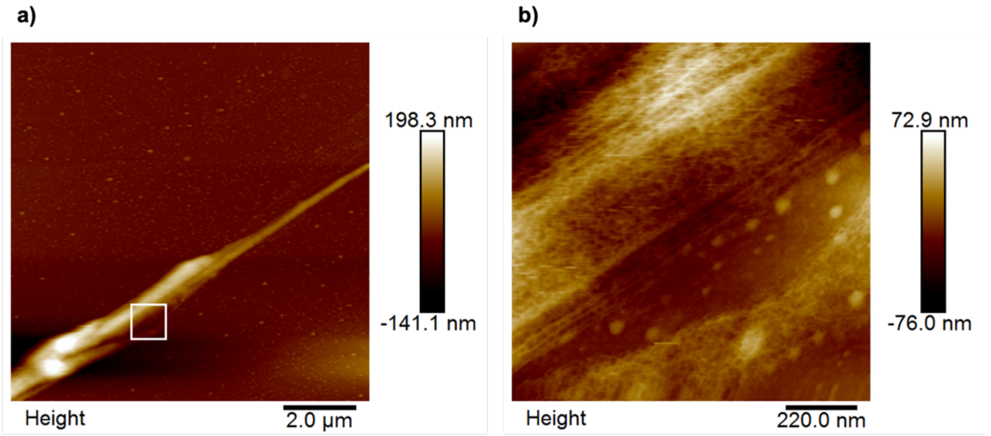


**Figure S6. Micron-scale fibres form from FUS condensates upon application of shear forces. (a,b)** AFM images of elongated structures when aged condensates ($t_a$=24 h) were deposited via manual deposition. A zoom of the structure in (a) is indicated by a white square. High-resolution imaging reveals a mesh-like network in these fibre-like structures (b).

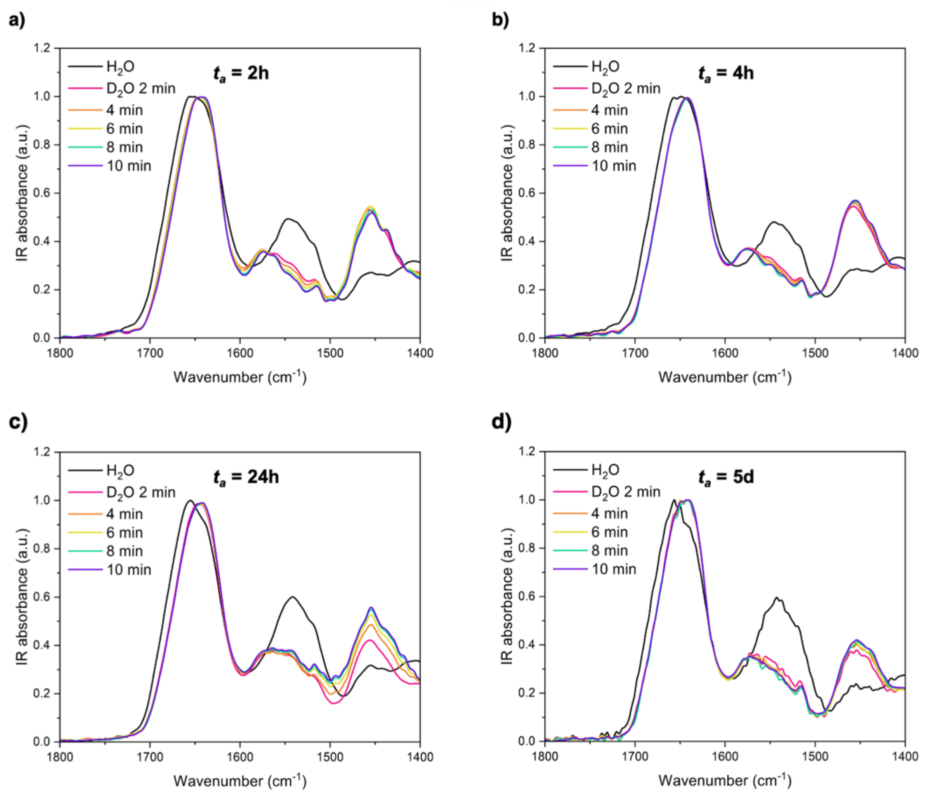


**Figure S7. HDX measurements on other time points. (a-d)** Spectra in $H_2O$ (black line) and $D_2O$ (rainbow lines) were plotted for $t_a$ = 2 h (a), 4 h (b), 24 h (c), and 5 d (d).

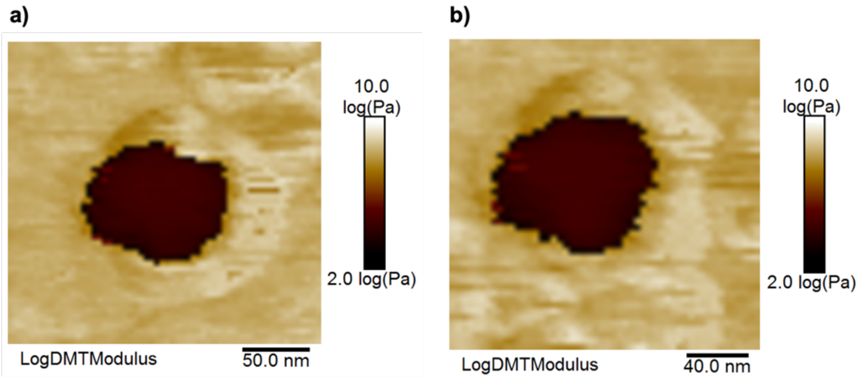


**Figure S8. Nanomechanical imaging repeated on the same condensate. (a,b)** Condensates were imaged (a), then re-imaged (b) to assess whether nanomechanical characterisation altered the morphology or nanomechanical properties. No detectable differences were observed when condensates were re-imaged.

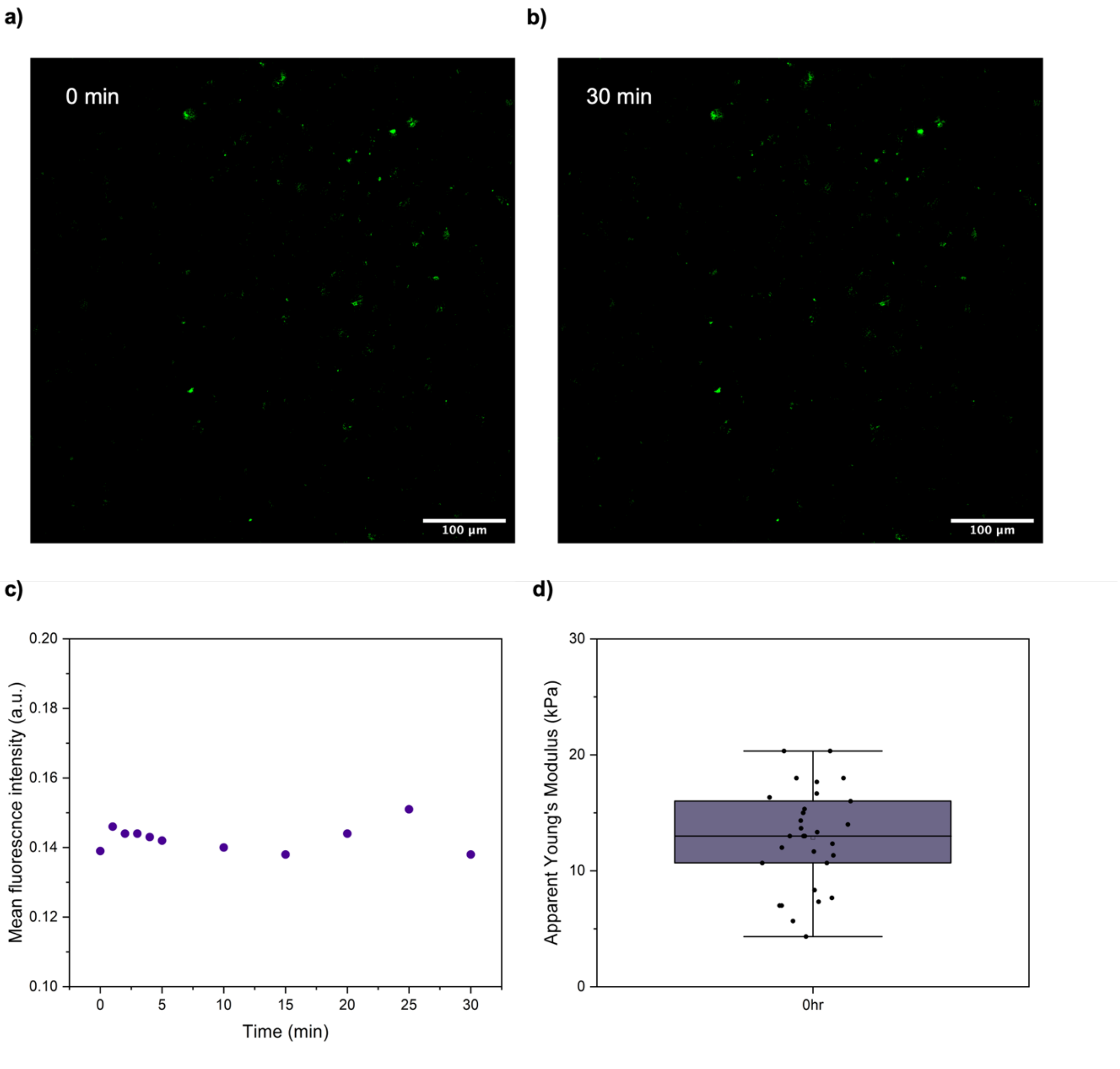


**Figure S9. Material properties of non-aged condensates. (a-b)** FUS condensates were deposited via microfluidic spray and imaged via confocal microscopy at 0 min (a) and after 30 min (b) after the addition of hexane-diol. **(c)** Average fluorescent intensity was plotted over time after addition of hexane-diol, and no change was observed, indicating condensates did not dissolve. **(d)** Apparent Young's modulus values of sample deposited via microfluidic spray at $t_a$ = 0 h are higher than that expected for a freshly-formed, liquid condensate.

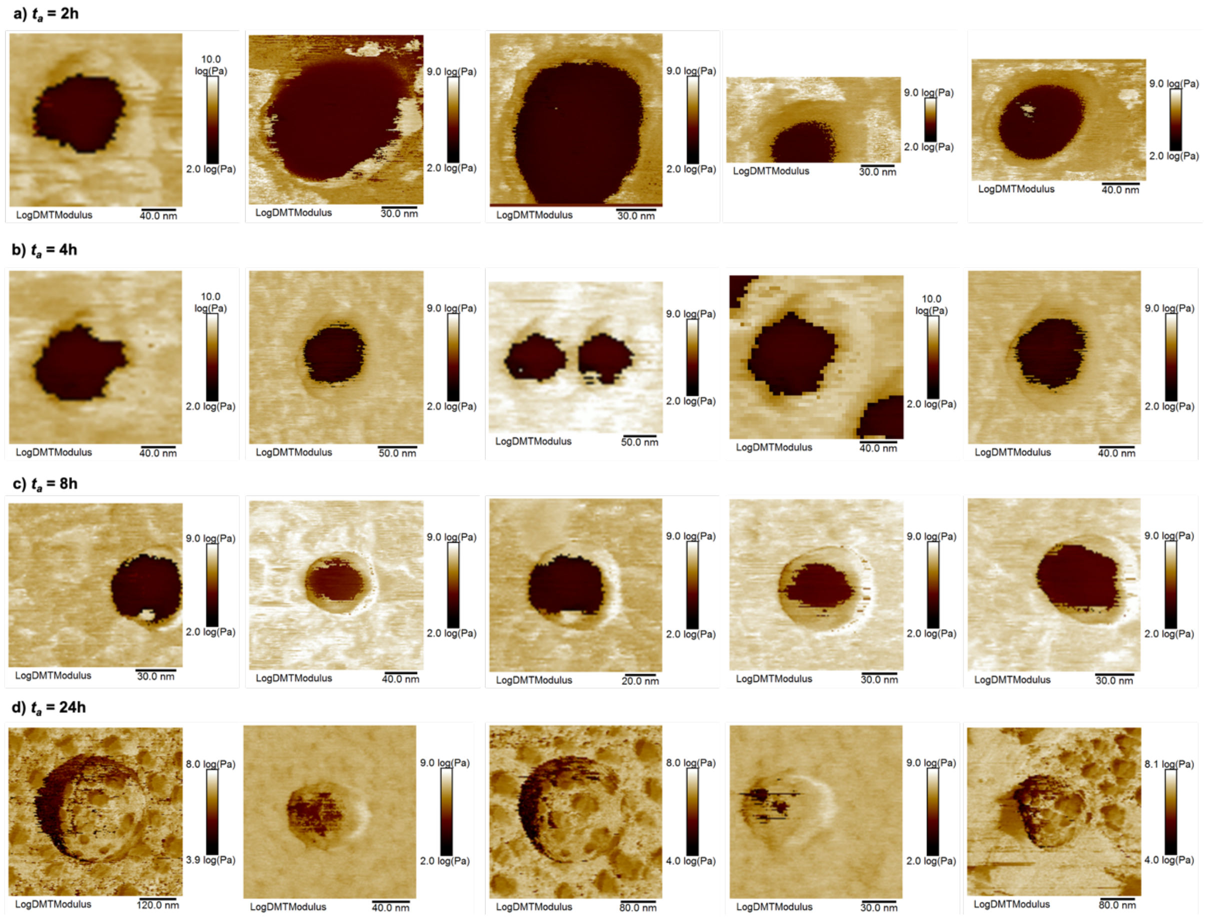


**Figure S10. Nanomechanical maps of condensates.** (a-d) Additional representative images, acquired via nanomechanical mapping, are shown of condensates as a function of $t_a$ = 2h (a), 4h (b), 8h (c) and 24h (d).

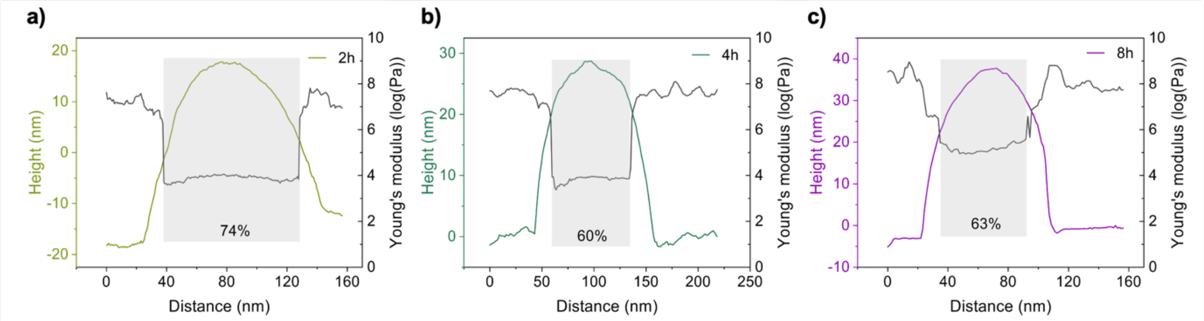


**Figure S11. Solid phase formation at condensate-solvent interfaces. (a-c)** Representative cross-sections of the height and Young's modulus for condensates at $t_a$=2 h (a), 4 h (b) and 8 h (c), taken at the centre of condensates. The percentage of the low elastic modulus phase for each condensate was calculated by considering the Young's modulus cross-section vs the condensate width (from the height profile). If the observed high elastic modulus phase emergence at condensate-solvent interfaces were solely due to the edge effect artefact, then the percentage of the low modulus phase would be smaller for softer condensates at $t_a$ = 2 h. However, the opposite is observed, with a decrease in the low modulus percentage at later aged condensates ($t_a$ = 4 and 8 h). We can also observe from these cross-sections that the high-modulus 'shell' is lower than the surface alone. This indicates that the observation of a high modulus, solid-like shell is not solely due to the edge effect artefact.

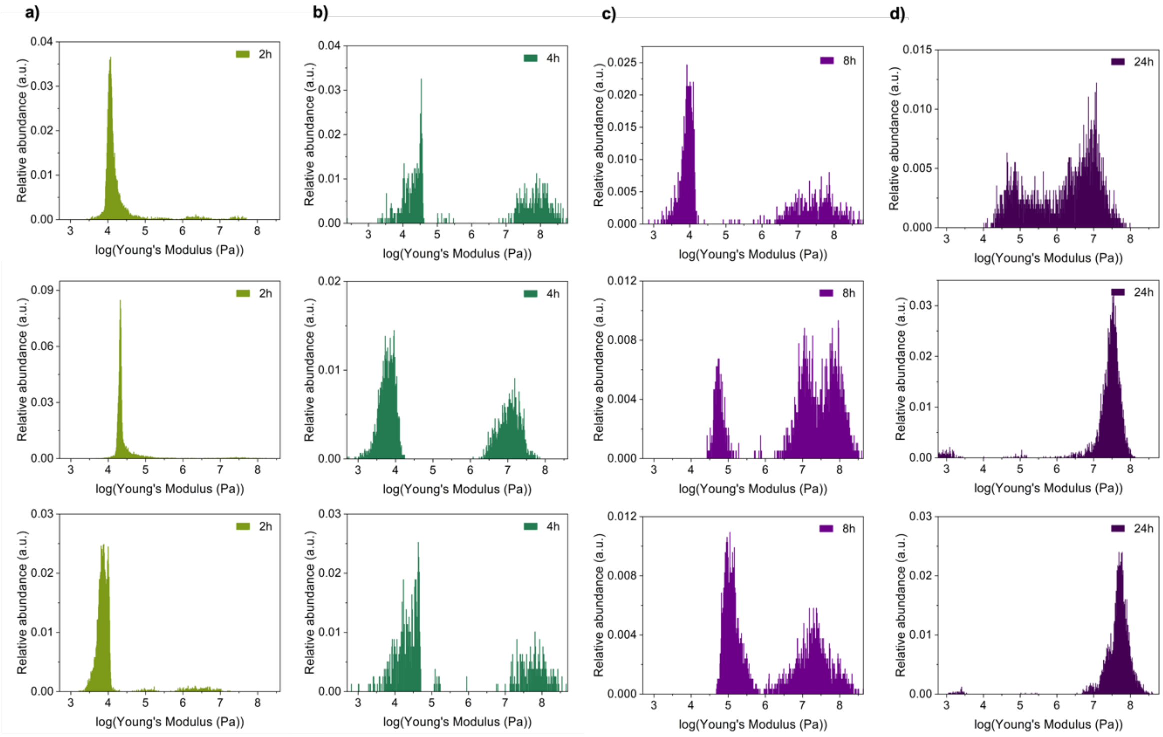


**Figure S12. Phase distributions within single condensates. (a-d)** Histograms of the Young's modulus distributions within single condensates are presented for $t_a$ = 2 h (a), 4 h (b), 8 h (c) and 24 h (d). These plots reveal the presence of distinct phases: a low elastic modulus phase, an intermediate elastic modulus phase, and a high elastic modulus phase.

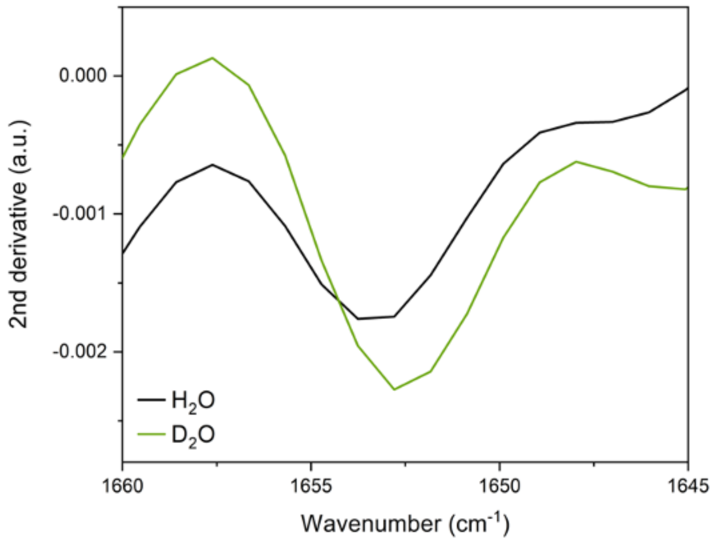


**Figure S13. α-helix absorption peak for late-aged samples in $H_2O$ and $D_2O$.** Spectra were acquired at $t_a$ = 8 h. The 2nd derivative is plotted in $H_2O$ (black) and $D_2O$ (green). The peak minimum in derivative plot in $H_2O$ is at 1653 $cm^{-1}$, and we observe a downward shift of 1 $cm^{-1}$ in $D_2O$. This is characteristic of α-helical secondary structure.

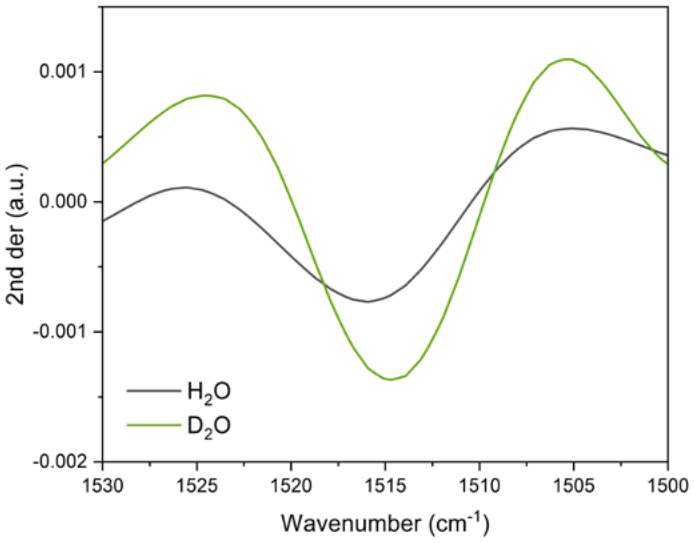


**Figure S14. Tyrosine absorption peak in $H_2O$ and $D_2O$.** Spectra were acquired at $t_a$ = 0 h. The 2nd derivative is plotted in $H_2O$ (black) and $D_2O$ (green). The peak minimum in derivative plot in $H_2O$ is at 1517 $cm^{-1}$, and we observe a downward shift of 4 $cm^{-1}$ in $D_2O$. This is characteristic of the tyrosine benzene ring C=C stretching vibration.

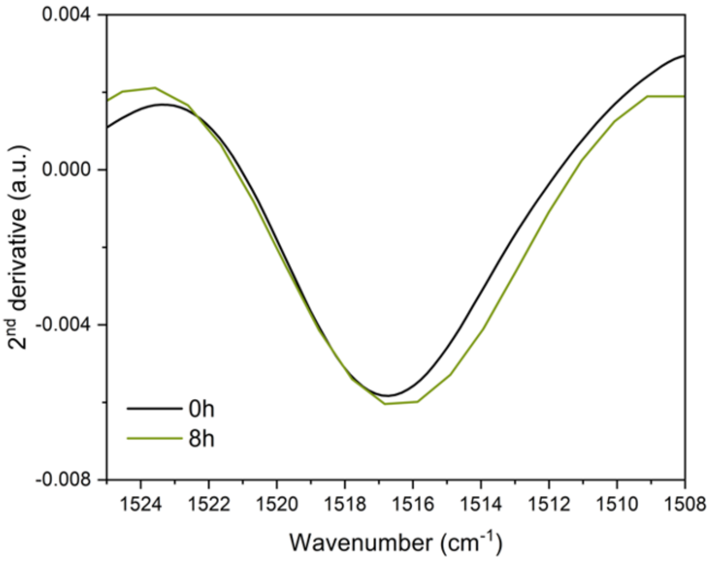


**Figure S15. Tyrosine peak position at $t_a$ = 8 h.** Comparison of the tyrosine peak position for condensates at $t_a$ = 0 and 8 h.
